# Ionotropic receptors in the turnip moth *Agrotis segetum* respond to repellent medium-chain fatty acids

**DOI:** 10.1101/2021.02.25.432965

**Authors:** Xiao-Qing Hou, Dan-Dan Zhang, Daniel Powell, Hong-Lei Wang, Martin N. Andersson, Christer Löfstedt

## Abstract

**Background:** In insects, airborne chemical signals are mainly detected by two receptor families, odorant receptors (ORs) and ionotropic receptors (IRs). Functions of ORs have been intensively investigated in Diptera and Lepidoptera, while the functions and evolution of the more ancient IR family remain largely unexplored beyond Diptera.

**Results:** Here, we identified a repertoire of 26 IRs from transcriptomes of female and male antennae, and ovipositors in the moth *Agrotis segetum*. We observed that a large clade formed by IR75p and IR75q expansions is closely related to the acid-sensing IRs identified in Diptera. We functionally assayed each of the five AsegIRs from this clade using *Xenopus* oocytes and found that two receptors responded to the tested ligands. AsegIR75p.1 responded to several compounds but hexanoic acid was revealed to be the primary ligand, and AsegIR75q.1 responded primarily to octanoic acid, and less so to nonanoic acid. It has been reported that the C_6_-C_10_ medium-chain fatty acids repel various insects including many drosophilids and mosquitos. Our GC-EAD recordings showed that C_6_-C_10_ medium-chain fatty acids elicited antennal responses of both sexes of *A. segetum*, while only octanoic acid had repellent effect to the moths in a behavioural assay. In addition, using fluorescence *in situ* hybridization, we demonstrated that the five IRs and their co-receptor AsegIR8a are not located in coeloconic sensilla as found in *Drosophila*, but in basiconic or trichoid sensilla.

**Conclusions:** Our results significantly expand the current knowledge of the insect IR family. Based on the functional data in combination with phylogenetic analysis, we propose that subfunctionalization after gene duplication plays an important role in the evolution of ligand specificities of the acid-sensing IRs in Lepidoptera.

## Introduction

Insects have evolved the ability to accurately sense environmental chemical signals, which play critical roles for their survival and reproduction (Sachse and Krieger, 2011; Leal, 2013). The peripheral process of chemoreception relies on receptors that interact with external molecular cues, triggering the transduction of chemical signals into electrical signals, which may ultimately result in a behavior (Hansson and Stensmyr, 2011; Fleischer et al., 2018). In insects, airborne chemicals are mainly detected by ligand-activated receptors from two families, namely the odorant receptors (ORs) (Clyne et al., 1999) and ionotropic receptors (IRs) (Benton et al., 2009). In *Drosophila*, these two receptor families are present in the dendritic membrane of the olfactory sensory neurons (OSNs) housed in different types of sensilla with distinct chemical preferences: ORs expressed predominantly in trichoid and basiconic sensilla detect, for example, pheromones and plant volatiles, including alcohols, aldehydes, esters and aromatics; IRs, mainly expressed in coeloconic sensilla, primarily respond to acids and amines (Yao et al., 2005; Benton et al., 2009; Abuin et al., 2011; Zhang et al., 2019).

IRs originated from the ionotropic glutamate receptor (iGluR) gene family and share similar structure and mechanism of action as iGluRs. However, the expression pattern and function of the IR family differ from the three well-studied iGluR families; kainate, NMDA (N-methyl-D-aspartate) and AMPA (α-amino-3-hydroxy-5-methyl-4-isoxazolepropionic acid) receptors that are expressed on the surface of synapses in animal nervous systems and play a role in signal transmission between neurons (Benton et al., 2009; Abuin et al., 2011). Insect IRs are present in different tissues and are more diverse in function. IRs were first identified in the model insect *Drosophila melanogaster* as odorant sensing receptors (Benton et al., 2009), and subsequent studies extended our knowledge on their functions to the involvement in taste (Koh et al., 2014; Stewart et al., 2015; Ahn et al., 2017; Chen and Amrein, 2017), temperature (Chen et al., 2015; Ni et al., 2016), humidity (Enjin et al., 2016; Knecht et al., 2016; 2017) and even auditory sensing (Senthilan et al., 2012). Similar to the ORs that are co-expressed with the conserved odorant receptor co-receptor (Orco) to form a functional ligand-gated ion channel (Larsson et al., 2004; Sato et al 2008; Wicher et al., 2008), co-expression in neurons is also observed among insect IRs. These receptors display more complex combinations since at least three IRs (IR8a, IR25a and IR76b) can function as co-receptors of specific tuning IRs. IR8a is normally co-expressed with IRs tuned to acids (Grosjean et al., 2011; Silbering et al., 2011; Rytz et al., 2013; Ai et al., 2013), while IR25a and IR76b are co-expressed with IRs responding to amines (Abuin et al., 2011; Hussain et al., 2018). In addition to the role in olfaction, IR25a was also reported as a co-receptor for IRs detecting temperature, humidity, and taste molecules (Ni et al., 2016; Enjin et al., 2016; Ganguly et al., 2017).

Based on their phylogeny and expression profiles, *Drosophila* IRs are classified into two groups: antennal IRs (A-IRs) and divergent IRs (D-IRs). A-IRs are highly conserved across insects and mainly expressed in the antennae, whereas the D-IRs have radiated to different extents in different species and are present in various tissues (Benton et al., 2009; Croset et al., 2010; Abuin et al., 2011; Rytz et al., 2013; Koh et al., 2014; Andersson et al., 2019; He et al., 2019). A number of the so-called Lepidoptera-specific IRs (LS-IRs) are absent in *Drosophila* and were originally identified in Lepidoptera (Olivier et al., 2011; Bengtsson et al., 2012; Poivet 2013; van Schooten et al., 2016; Liu et al., 2018; Zhu et al., 2018a). Despite the name, some of them are also present in Trichoptera and mosquitos with overall high support values, although in some cases the orthology remains somewhat ambiguous (Yuvaraj et al., 2018; Yin et al., 2020). The LS-IRs do not represent a single monophyletic IR subfamily, because they are scattered across the phylogenetic tree, with some (IR1, IR75p, IR75q and IR87a) located within the A-IR while others (IR7d, IR100b-j and IR143) nested within the D-IR clade, and also include a putative pseudogene (IR2) (Liu et al., 2018; Zhu et al., 2018a). The large IR75p and IR75q expansions are of particular interest due to their close relationship with the characterized acid-sensing IRs (Liu et al., 2018; Zhu et al., 2018a).

Over the past decade, much progress has been made in elucidating the functions of IRs in *Drosophila* and a few mosquito species. Although this major gene family spans across protostomes (Croset et al., 2010; Eyun et al., 2017), the functional information of IRs beyond dipterans is still limited, restricting our knowledge concerning the evolution and specialization of this ancient gene family. To our knowledge, only two IRs from the parasitoid wasp *Microplitis mediator* have been functionally investigated (Shan et al., 2019). The second largest insect order, Lepidoptera, rely on their remarkable olfactory abilities to sense the external environment. However, the function of IRs in Lepidoptera remains largely unexplored, although recent studies in *Manduca sexta* and *Mythimna separata* suggested a role for the co-receptor IR8a in acid-sensing (Zhang et al., 2019; Tang et al., 2020). In the present study, we report the first functional characterization of IRs in Lepidoptera. We show that two IRs from the turnip moth *Agrotis segetum* respond to medium-chain fatty acids and that these fatty acids elicit antennal responses, with octanoic acid being significantly repellent to moths in an olfactory bioassay. Further, we find that in this moth the acid-detecting IRs are expressed in basiconic or trichoid sensilla, but not in coeloconic sensilla as was found in *Drosophila*. Based on our phylogenetic analysis and receptor characterization, we suggest that subfunctionalization occurred in Lepidoptera after a gene duplication event in the IR75p/q clade.

## Results

### *Characterization of the* A. *segetum IR repertoire from antennal and ovipositor transcriptomes*

Studies on *M. sexta* have shown that in addition to the antennae, the ovipositor plays an important chemosensory role (Klinner et al., 2016). Hence, in the present study we profiled replicated transcriptomes of the antennae (N=4 for both sexes) and ovipositors (N=2) from *A. segetum* (‘Aseg’) by mRNA sequencing (RNA-Seq) for an exhaustive search of IRs involved in chemosensation. This produced a total of over 50 gigabases of sequence data with the yield of each sample reaching over 10 million paired-end (100 bp) reads. Clean reads from all 10 samples were assembled together to produce a transcriptome comprised of 83,645 non-redundant transcripts having a mean length of 976 bp and an N_50_ of 1,788 bp.

In total, 26 AsegIR genes were identified, including the three co-receptors, IR8a, IR25a and IR76b. Of the 26 assembled IR transcripts, 23 encoded full-length genes. We followed the nomenclature previously used to name lepidopteran IRs (Liu et al., 2018). The raw sequence reads have been deposited in the SRA database at NCBI under the Bioproject accession number PRJNA707654.

### The IR75p and IR75q lineages grouped with acid-sensing IRs

We performed phylogenetic analyses of AsegIRs along with the IR sequences from *Bombyx mori* (Bombycidae), *Plutella xylostella* (Plutellidae), *H. armigera* (Noctuidae), *D. melanogaster* and *Anopheles gambiae* (both Diptera) (Fig. 1). The two IR classes, A-IRs and D-IRs, identified in previous studies (Croset et al., 2010; Liu et al., 2018; Zhu et al., 2018a) can be recognized in the tree. A single AsegIR orthologue from all the 18 monophyletic groups of conserved A-IRs (IR1.1, IR1.2, IR21a, IR31a, IR40a, IR41a, IR60a, IR64a, IR68a, IR75d, IR75p, IR75p.1, IR75p.2, IR75q.1, IR75q.2, IR76b, IR87a, IR93a) were identified. Fewer AsegIRs were identified in the D-IR class, which was expected because D-IRs are typically expressed in tissues other than antennae and ovipositors, and presumably mediate non-olfactory behaviors. The large expansions of IR7d, IR75p and IR75q have been evident in almost all the reported antennal transcriptomes of Lepidoptera, and these IRs typically have relatively high levels of gene expression. The expression of IR75p and IR75q genes is predominantly restricted to the adult antennae, whereas IR7d genes seem to be generally distributed in different body parts of adults and larvae (van Schooten et al., 2016; Liu et al., 2018; Zhu et al., 2018a). Notably, the lineage expansions IR75p and IR75q, along with IR1.1 and IR1.2, grouped within a larger cluster together with IR75a, IR64a and IR31a, which have been reported as acid-sensing IRs in *Drosophia* and mosquitos (Abuin et al., 2011; Ai et al., 2013; Silbering et al., 2011), although IR75d in the neighbouring expansion responded to an amine (pyrrolidine) in *Drosophila* (Silbering et al., 2011).

**Fig. 1.**
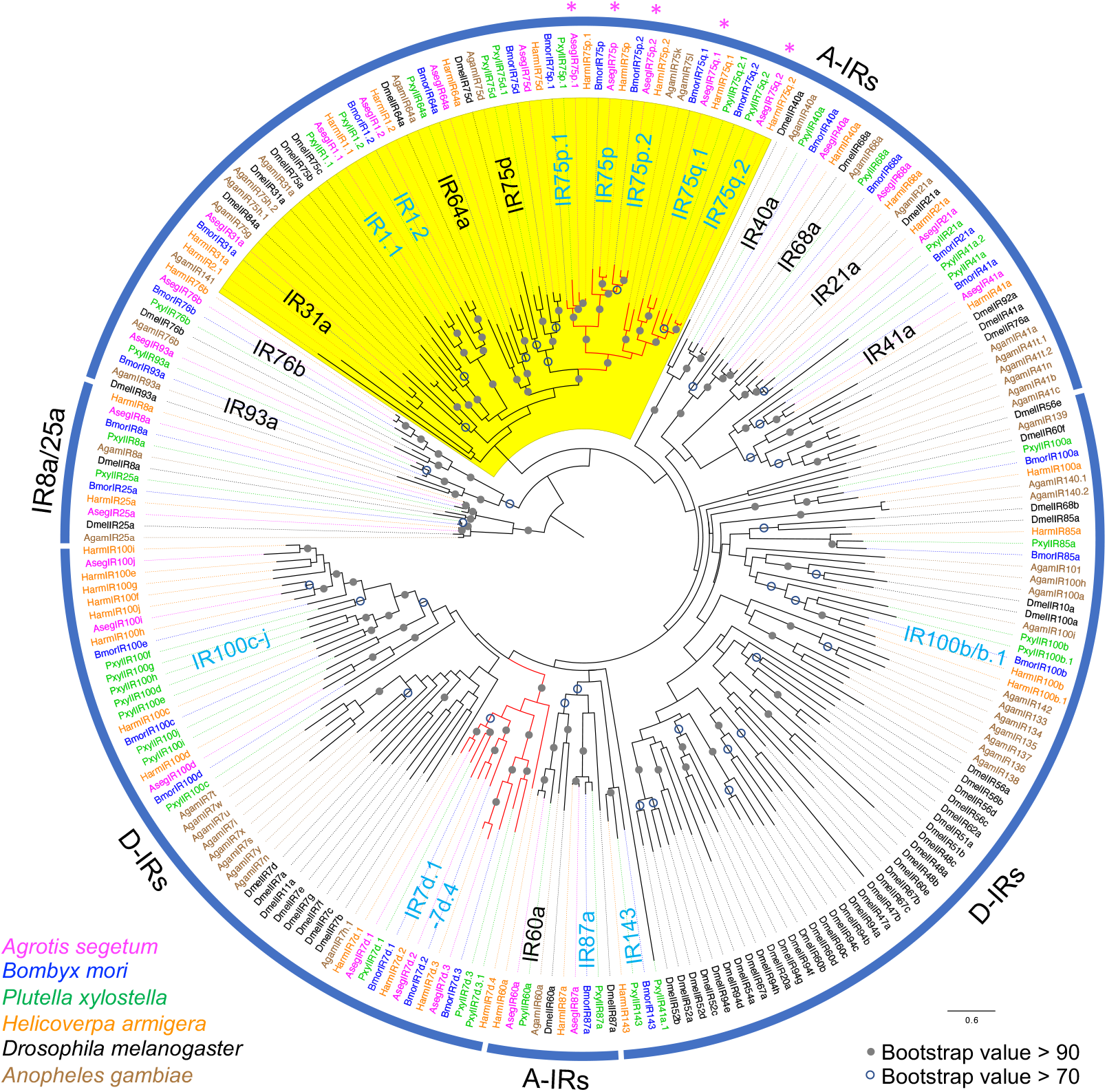
Phylogeny of Lepidoptera IRs. The maximum-likelihood phylogenetic tree is based on a protein sequence alignment of ionotropic receptors (IRs) from *A. segetum*, *B. mori* (Croset et al., 2010), *P. xylostella* (Liu et al., 2018), *H. armigera* (Liu et al., 2018), *D. melanogaster* (Benton et al., 2009; Croset et al., 2010) and *A. gambiae* (Croset et al., 2010; Liu et al., 2010). The tree was rooted with the lineage of IR8a and IR25a receptors. The IRs from different species are color coded as indicated in the figure. The monophyletic IR groups are noted on the tree, with the light blue color indicating “Lepidoptera-specific IR groups”. The large IR7d, IR75p and IR75q expansions are indicated by red branches, and the putative “expanded acid-sensing cluster” is highlighted in yellow with the exception of IR75d which responded to an amine in *Drosophila*. The five IR75p and IR75q receptors functionally assayed in this study are marked with asterisks. Bootstrap support (100 replicates) values over 0.7 are shown at corresponding nodes.

### AsegIRs in 75p/q expansions were predominantly expressed in adult antennae

The expression levels of the 26 AsegIRs were normalized across sequencing libraries using Transcripts Per Million (TPM) scaling factor (Li and Dewey, 2011). Surprisingly, one of the LS-IRs, AsegIR7d.3, was found to be the most abundantly expressed IR, with much higher levels of expression than that of the three co-receptors, and with higher expression in ovipositors than in both male and female antennae. Among the three co-receptors, AsegIR25a showed the highest expression, with higher TPM values compared to AsegIR8a and AsegIR76b. AsegIR25a and AsegIR76b had non-biased expression in the antennae of both sexes, while AsegIR8a showed female-biased expression. AsegIR75q.1 and AsegIR75q.2 were identified from a large polycistronic transcript with comparatively high expression in both male and female antennae. The other three IRs in the IR75 clade, i.e., IR75p, IR75p.2 and especially IR75p.1, were expressed at much lower levels (Fig. 2A).

**Fig. 2.**
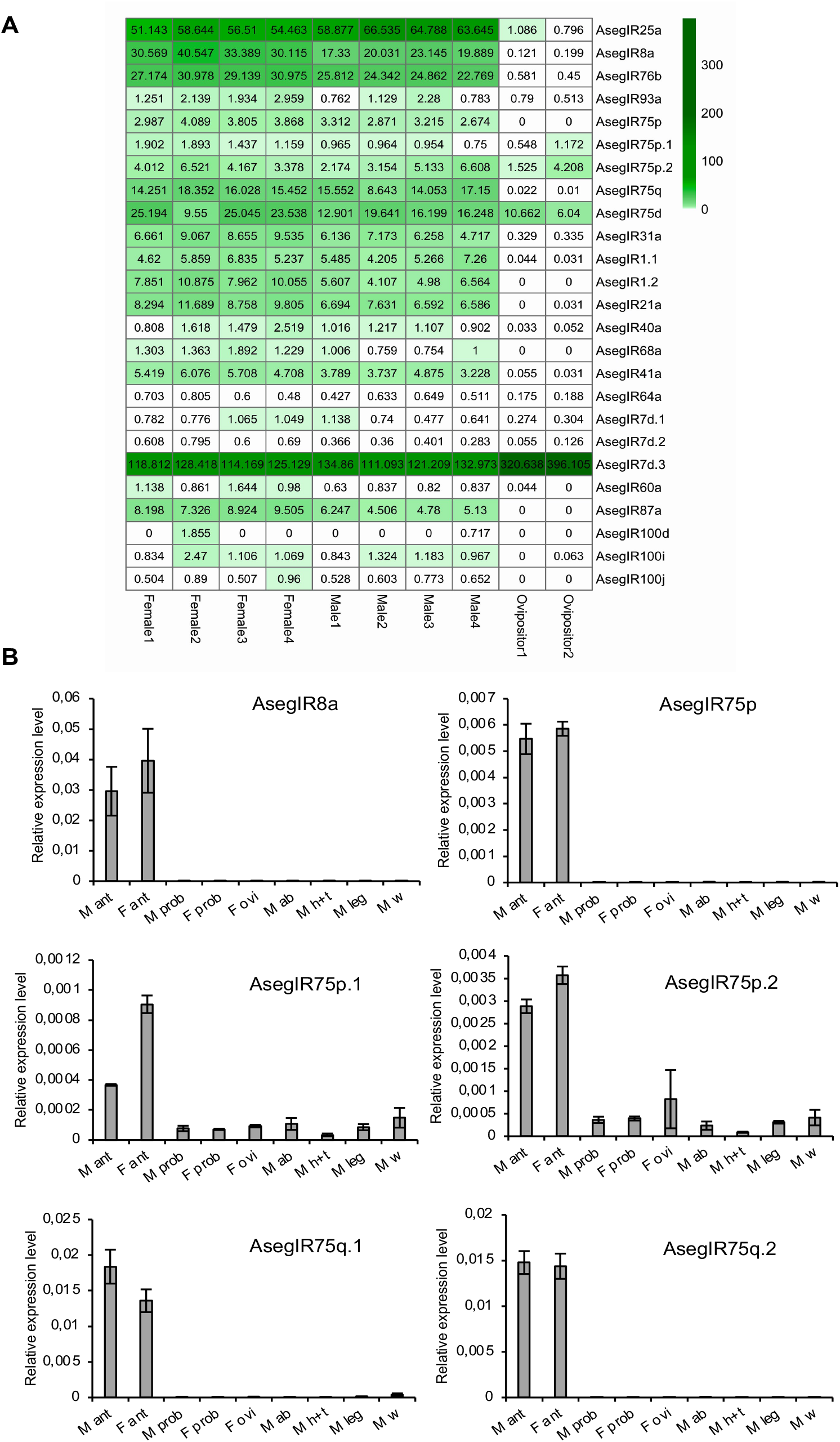
The expression profiles of AsegIRs. **(A)** Heatmap with TPM value (Transcripts Per Million) showing the expression levels of the AsegIRs in different female and male antennal samples, as well as ovipositor samples. **(B)** The expression of AsegIRs from IR75p and IR75q lineages and the co-receptor IR8a in different tissues and sexes. M, male; F female; ant, antennae; prob, proboscis; ovi, ovipositors; ab, abdomens; h+t, head and thorax (without antennae); w, wings. As AsegIR75q.1 and AsegIR75q.2 are located on the same transcript, it is not possible to distinguish the TPM value for each of them from the transcriptome data, whereas the qPCR results demonstrated that both were more abundant than the other three AsegIRs in the IR75p/q expansion.

Quantitative PCR was conducted to further investigate the expression pattern of the AsegIRs from the IR75p/q expansions and their putative co-receptor IR8a in various tissues of both sexes, including chemosensory organs (antennae and proboscises) of both sexes, female ovipositor, as well as male abdomen, head and thorax (without antennae), legs and wings. GADPH (glyceraldehyde 3-phosphate dehydrogenase) and RPS3 (ribosomal protein S3) were used as reference genes. The results showed that AsegIRs from the IR75p/q expansions and AsegIR8a were expressed predominantly in adult antennae of both sexes, although IR75p.1 and IR75p.2 showed a broader tissue distribution, being weakly expressed in almost all assayed non-chemosensory tissues (Fig. 2B). Consistent with the results of the transcriptome analysis, AsegIR8a had higher expression level in female antennae than in male antennae; AsegIR75q.1 and AsegIR75q.2 were much more abundantly expressed than the other IR genes in the expansion, while IR75p.1 had a lower expression level and was female-biased (Fig. 2B).

### AsegIRs are expressed in basiconic or trichoid sensilla

In *Drosophila*, IRs are mainly located in the antennal coeloconic sensilla (Benton et al., 2009; Rytz et al., 2013), whereas it was previously unknown which sensillum type the IRs in moths are associated with. To address this question, we first studied the morphology and distribution of the sensilla on *A. segetum* antennae by scanning electron microscopy (SEM). The antennae of *A. segetum* are sexually dimorphic, with setaceous antennae in females and plumose antennae in males (Fig. 3A–D). There is one pair of branches on most of the segments of male antennae, whereas the segments towards the tip of the antennae are unbranched (Fig. 3B and 3D). The SEM images indicated that the trichoid sensillum was the most abundant type on both female and male antennae and it was the only sensillum type present on the branches of male antennae (Fig. 3E, 3F and 3H). Only two or three coeloconic sensilla were observed on each segment of the male antennae, including the unbranched segments and the stem parts of the branched segments, but not on the branches; they were more abundant (six observed) on each segment of the female antennae (Fig. 3E and 3F). Basiconic sensilla were present throughout the entire female antennae and male antennae (except for the branches) but in lower abundance compared to trichoid sensilla in both sexes (Fig. 3E and 3F).

**Fig. 3.**
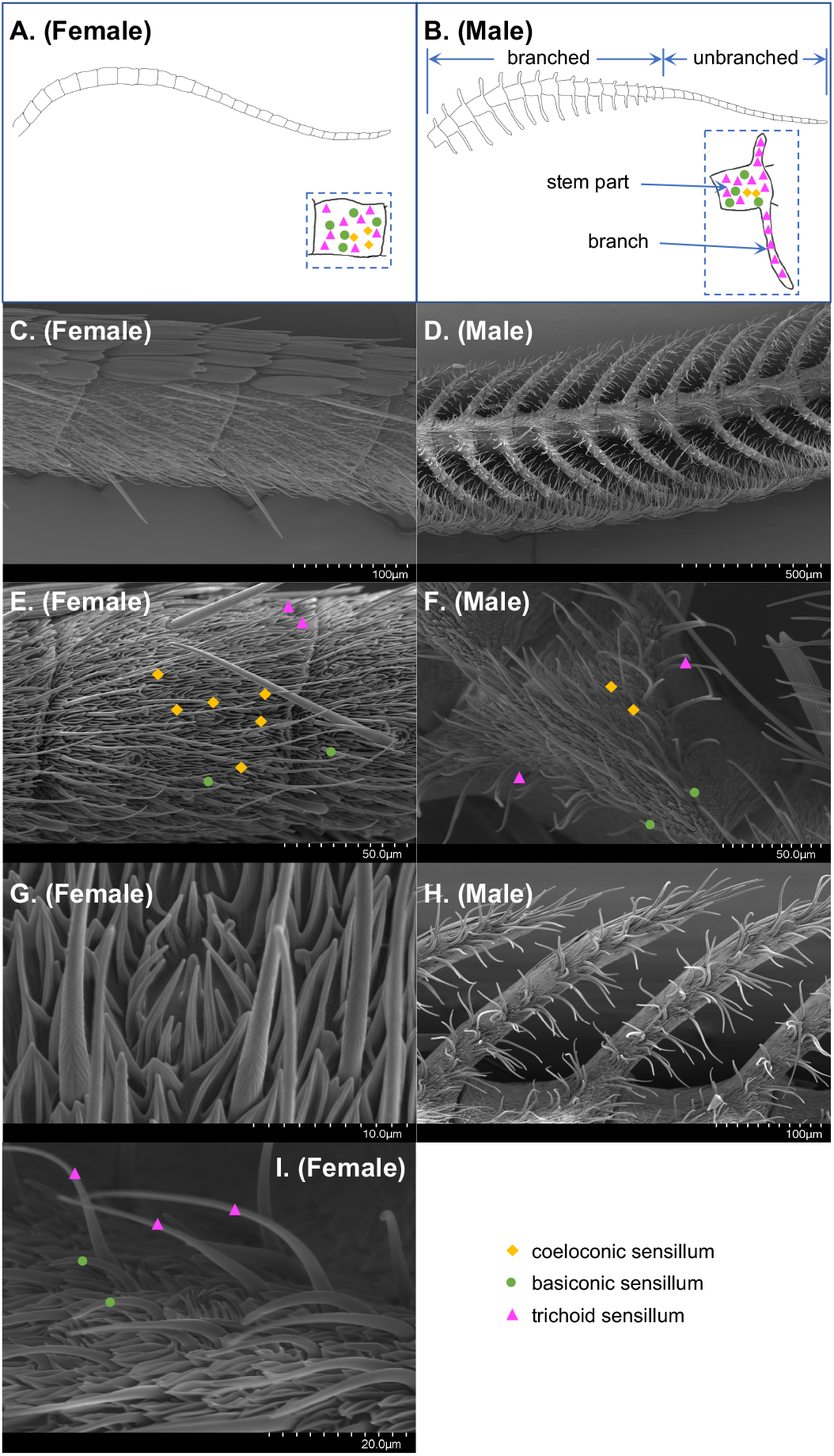
Scanning electron micrographs of *A. segetum* antennae. **(A)**-**(B)** Schematic diagrams and **(C**)-**(D)** overall SEM images showing the sexually dimorphic structures of female and male antennae of *A. segetum* moths. The distribution of coeloconic sensilla (yellow diamonds), trichoid sensilla (magenta triangles) and basiconic sensilla (green circles) are illustrated in the enlarged view of segments in the dashed frame. The stem part of male antennae, the branches and unbranched part are labelled in the figure. **(E**)-**(F)** Antennal segments showing representative coeloconic, trichoid, and basiconic sensilla. **(G)** Close-up view of a coeloconic sensillum. **(H)** Coeloconic sensilla are absent on the branches of male antennae. **(I)** Close-up view of trichoid sensilla and basiconic sensilla. The labelled number in the scales represents the length between the first and last marks, and the distance between two marks represents a tenth of the labelled number.

To further explore the distribution of AsegIR75p/q and their putative co-receptor AsegIR8a on female and male antennae and to confirm their co-expression pattern, we performed two-color fluorescence *in situ* hybridization (FISH) with digoxigenin (DIG)-labelled antisense RNA-probes for the five AsegIRs and biotin-labelled antisense probes for AsegIR8a. Corresponding sense probes were used as control. On female antennae, AsegIR8a was abundantly expressed across the whole segment, while signals of the five AsegIRs were much fewer (Fig. 4 and Fig. 5). On the male antennae, AsegIR8a and the five AsegIRs were predominantly expressed at the branched segments but not the unbranched segments toward the tip of the antennae (Fig. 4, Fig. 5 and Fig. S1). They were not expressed on the stretches of the branches (Fig. S1) but distributed at the bases of the branches and at the stem parts (Fig. 4 and Fig. 5). Fewer signals of the five AsegIRs were detected compared to AsegIR8a signals, consistent with their expression levels. The two-color FISH confirmed that the five AsegIRs were indeed co-expressed with AsegIR8a in both sexes (Fig. 4 and Fig. 5). Intriguingly, all the five AsegIRs as well as AsegIR8a were in both sexes not expressed in coeloconic sensilla on the antennae as in *Drosophila* (Fig. 4, Fig. 5 and Fig. S1). On the female antennae, they seemed to be expressed in the OSNs located in basiconic sensilla. On the male antennae, however, because the bases of the branches have high density of hair-like sensilla, we were not able to distinguish from the FISH images whether the IRs were expressed under basiconic sensilla or trichoid sensilla.

**Fig. 4.**
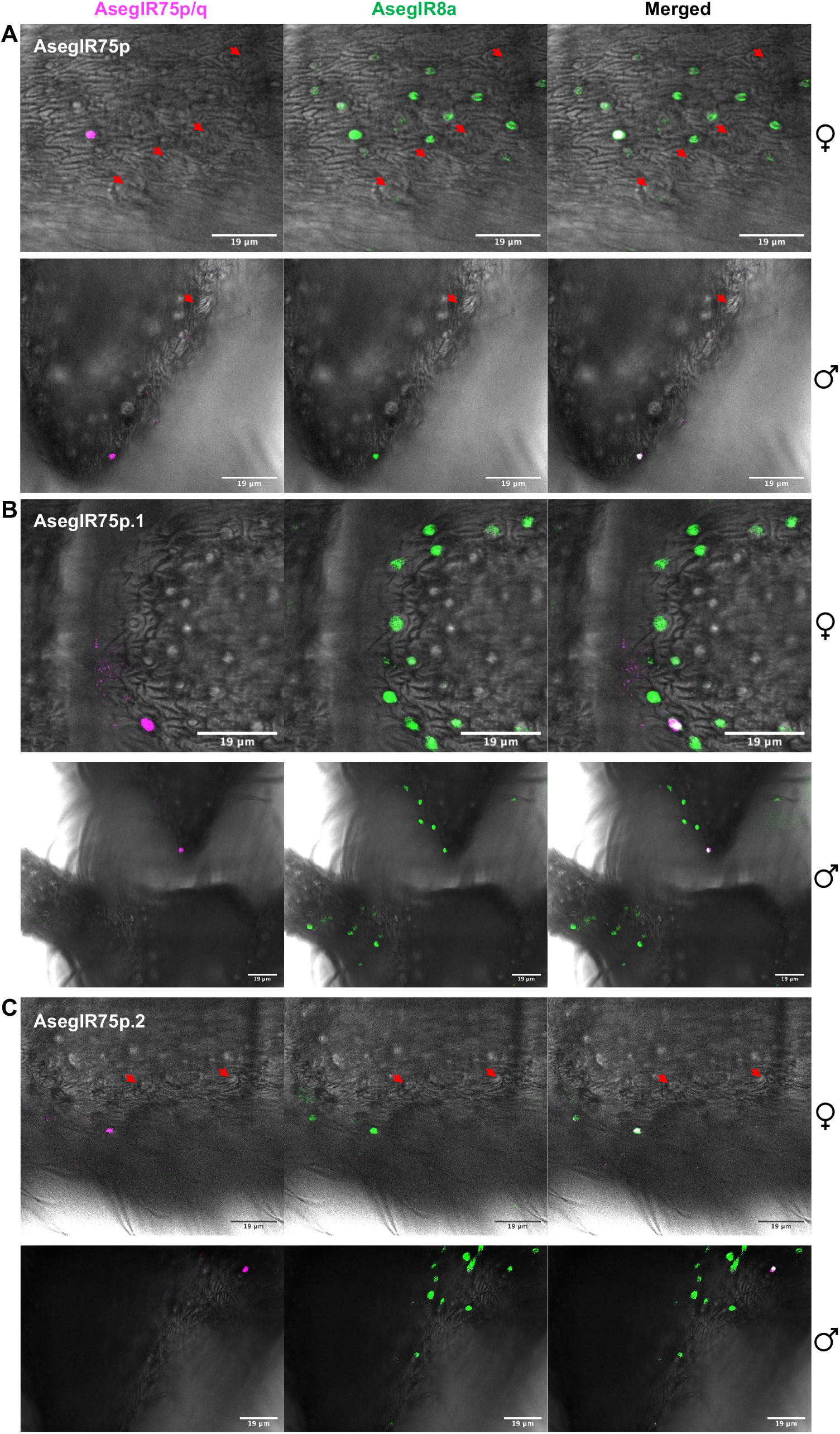
Whole-mount fluorescence *in situ* hybridization (WM-FISH) showing the co-expression of AsegIR8a with AsegIR75p (A), AsegIR75p.1 (B) and AsegIR75p.2 (C) on adult female and male antennae. Neurons expressing AsegIR8a are indicated in green, AsegIR75p/p.1/p.2 signals are indicated in magenta, and the merged signals are shown in white. Red arrowheads point to the location of coeloconic sensilla. The moths used in WM-FISH were 3-5 days old after eclosion and unmated.

**Fig. 5.**
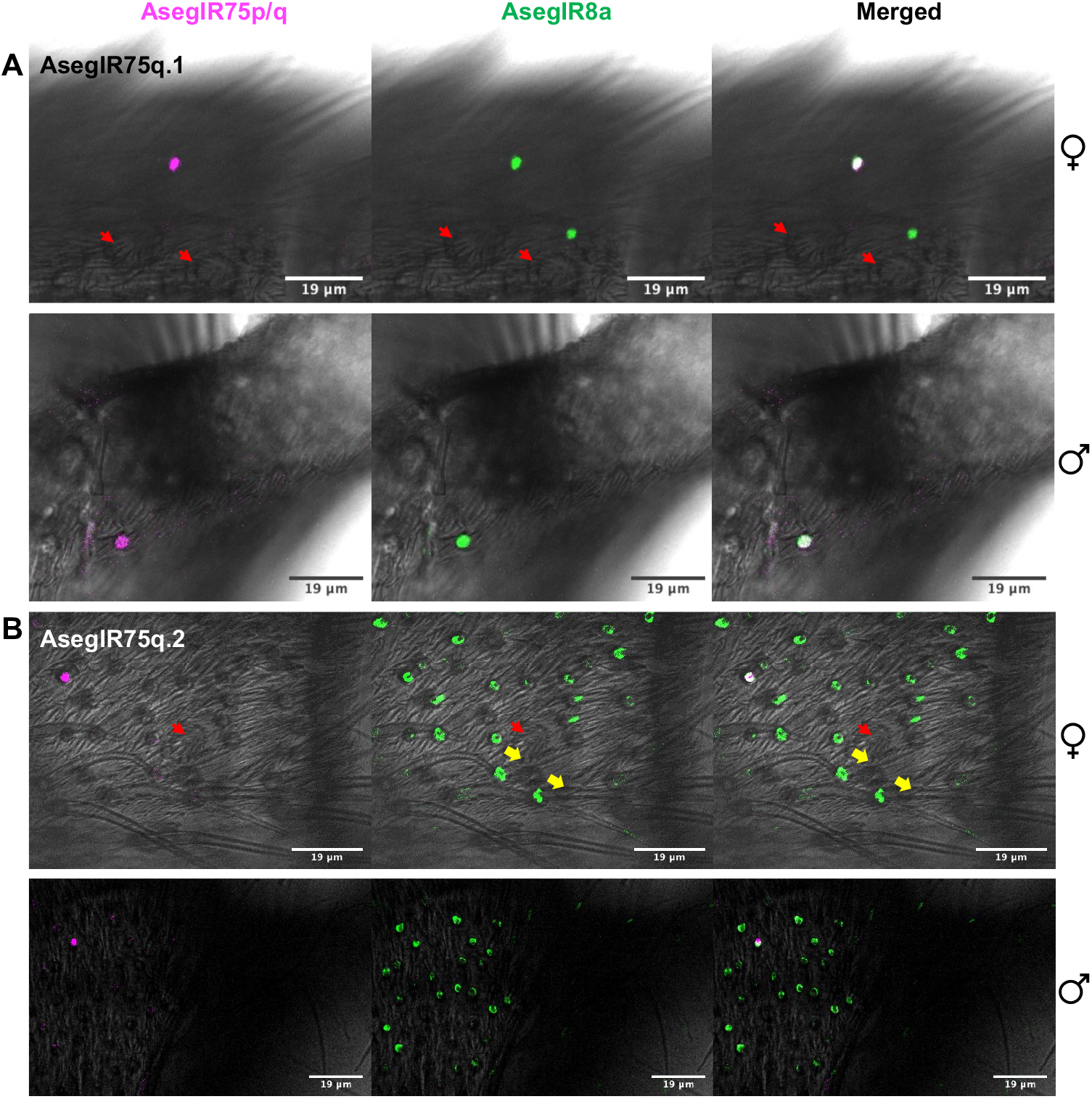
Whole-mount fluorescence *in situ* hybridization (WM-FISH) showing the co-expression of AsegIR8a with AsegIR75q.1 (A) and AsegIR75q.2 (B) on adult female and male antennae. Cell bodies of neurons expressing AsegIR8a are indicated in green, AsegIR75q.1/q.2 signals are indicated in magenta, and the merged signals are shown in white. Red arrowheads point to the location of coeloconic sensilla, while the yellow arrows point to the basiconic or trichoid sensilla with AsegIR expressing cell bodies below. The moths used in WM-FISH were 3-5 days old after eclosion and unmated.

### IR75p.1 and IR75q.1 responded to medium-chain fatty acids in Xenopus oocytes

The five AsegIRs from the IR75p/q expansions were each co-expressed with AsegIR8a in *Xenopus* oocytes, and screened against a panel of 46 compounds, most of which are common plant volatiles, including acids, aldehydes and alcohols. Oocytes co-expressing AsegIR75p.1/AsegIR8a showed responses to acids and aldehydes with exclusively C_5_-C_7_ straight chains at a concentration of 100 μM. For the alcohol ligands, only C_6_ unsaturated compounds elicited responses. The strongest response was elicited by hexanoic acid, followed by (*Z*)-3-hexenol, (*E*)-2-hexenal and hexanal (Fig. 6A and B). Dose-response trials indicated a higher sensitivity of AsegIR75p.1 to hexanoic acid compared to the other active ligands (Fig. 6B). Oocytes co-expressing AsegIR75q.1/AsegIR8a showed a strong response to octanoic acid and a weaker response to nonanoic acid at a concentration of 100 μM. This receptor also responded slightly to heptanoic acid and octanal, with response magnitudes much lower than that to the primary ligand (Fig. 6A and C). The responses to octanoic acid and nonanoic acid were dose-dependent, with a threshold concentration at 1 μM (Fig. 6C). The oocytes injected with the cRNAs of AsegIR75q.1, AsegIR75p.1 and AsegIR8a alone did not show any response to the tested compounds. In addition, the other three receptors in the same expansions, AsegIR75p, AsegIR75p.2 and AsegIR75q.2, did not respond to any of the tested compounds when co-expressed with AsegIR8a in oocytes.

**Fig. 6.**
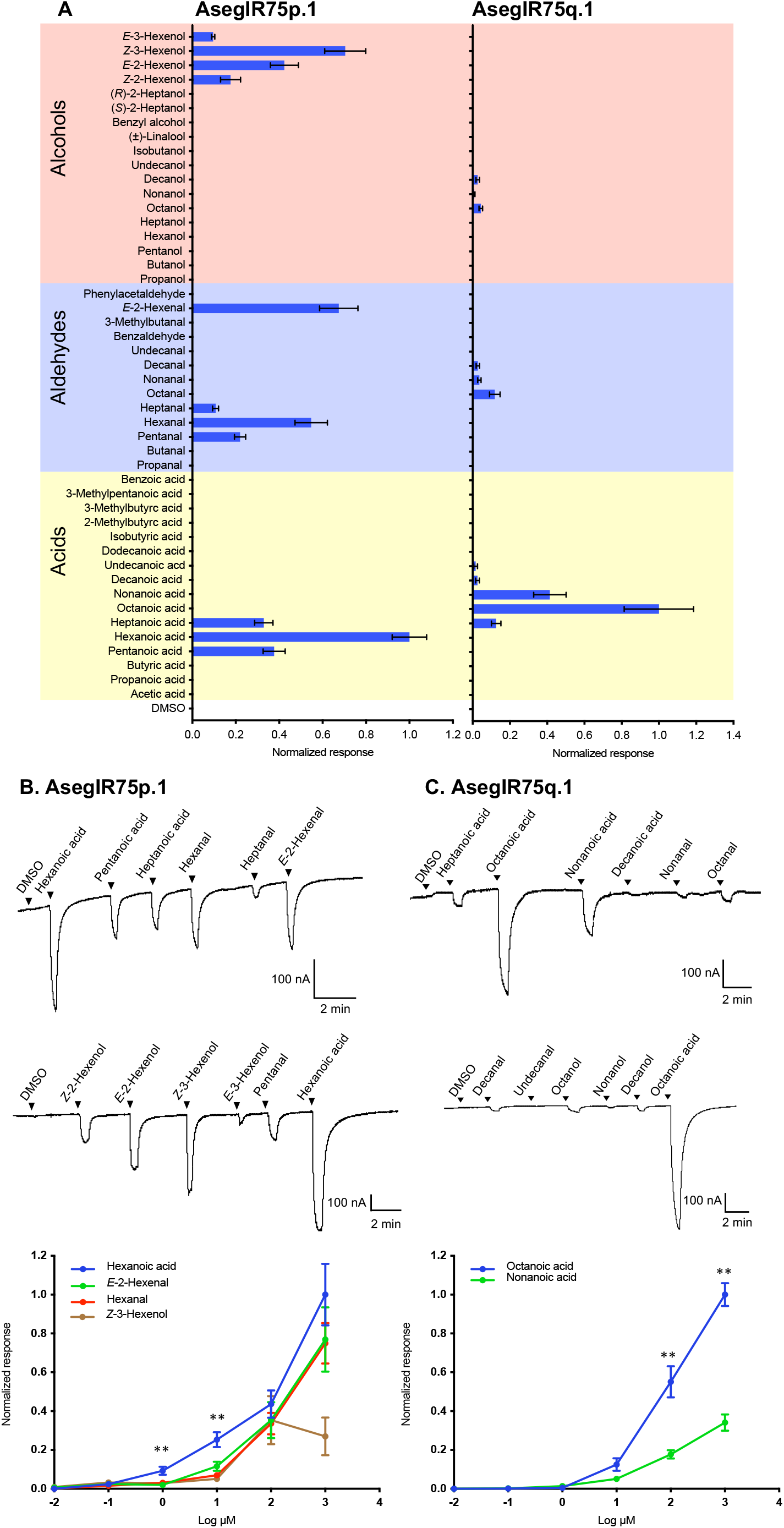
AsegIR75p.1 and AsegIR75q.1 respond to medium-chain fatty acids in *Xenopus* oocytes. **(A)** Response profiles of AsegIR75p.1 (left) and AsegIR75q.1 (right) to the full odor panel. Response magnitudes were normalized to the average response of primary ligand (N ≥4). Error bars indicate the SE. **(B)** and **(C)** Upper panel: Representative current traces of oocytes upon successive exposures to 100 μM stimuli. Each compound was applied at the time indicated by the arrowheads for 20 s. Lower panel: Dose-dependent responses, showing values normalized to the average response of the most active compound at 100 μM (N ≥4 for each ligand).

### Medium-chain fatty acids elicit antennal responses

Due to the responses of AsegIR75p.1 and AsegIR75q.1 to medium-chain fatty acids (C_6_-C_10_) in the oocytes, we further tested whether these acids could elicit antennal responses in the moths by gas chromatography coupled to electroantennographic detection (GC-EAD). Both female and male antennae showed clear responses to the tested acids (Fig. 7A). The responses to decanoic acid were smaller in both sexes, while the response magnitudes to the other tested acids were similar. Male moths seemed to have larger responses to hexanoic acid and octanoic acid than female moths, but the responses to the other tested acids were not significantly different between the two sexes (Fig. 7B).

**Fig. 7.**
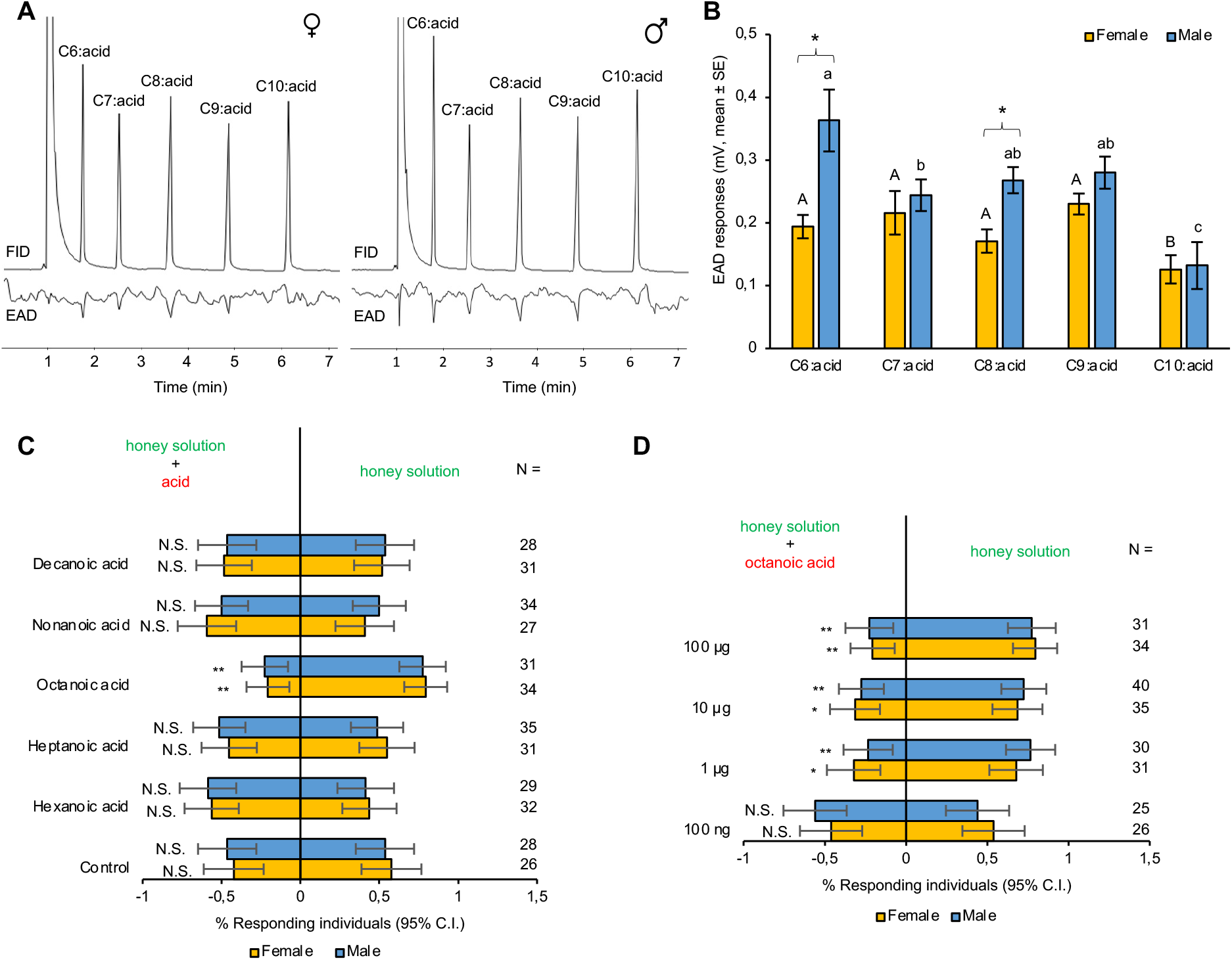
Antennal and behavioral responses of *A. segetum* to medium-chain fatty acids. **(A)** Representative electrophysiological responses of male and female *A. segetum* antennae to medium-chain fatty acids. Top: GC trace of the saturated C_6_-C_10_ acids abbreviated as 6:0, 7:0, 8:0, 9:0 and 10:0 (FID). Bottom: antennal responses to corresponding compounds (EAD). **(B)** GC-EAD responses of male and female *A. segetum* to C_6_-C_10_ acids. Values are amplitude of EAD responses (mV, mean ± SE) after being normalized by corresponding FID peak abundance. Statistically significant differences between stimuli within each sex are indicated by letters (one-way-ANOVA followed by an LSD test at 0.05 level). Differences between males and females responding to the same stimulus are indicated by asterisks after Student’s t-Test at p < 0.05 level. N=6 for female, N=4 for male. **(C)** Behavioral responses of female and male turnip moths in a Y-tube olfactometer to saturated fatty acids (with 6-10 carbon atoms) at the 100 μg dose, and **(D)** to octanoic acid at a series of doses from 100 ng to 100 μg. N value for each dual choice is shown on the right. Error bars indicate 95% confidence interval. P-values are based on chi-square tests: N.S., P>0.05; *, P<0.05; **, P<0.01.

### *Octanoic acid is behaviorally repellent to* A. *segetum*

In order to reveal the biological significance of the observed IR responses to fatty acids, we performed two-choice Y-tube bioassays. In an initial experiment, we tested whether the odor of honey solution (10%) was attractive to the adult moths. Among 51 individuals (29 females and 22 males), most of *A. segetum* from both sexes (female: χ^2^=12.45, d.f.=1, p= 0.0004; male: χ^2^=4.545, d.f.=1, p=0.033) were attracted to the honey solution and only two males chose water. In another control experiment, cotton balls with 10% honey solution were placed in each of the two arms of the Y-tube and were equally attractive to both female and male *A. segetum* (Fig. 7C), demonstrating no side-preference. In the treatment groups, individual female and male moths were given the choice between 10% honey solution and 10% honey solution loaded with one of the C_6_-C_10_ fatty acids at the dose of 100 μg. Our results indicated that among the five tested acids, only octanoic acid showed a significant repellent effect on both sexes, with significantly fewer individuals choosing the honey solution with octanoic acid compared to honey solution alone (female:χ^2^=11.7647, d.f.=1, p=0.0006; male: χ^2^=9.3226, d.f.=1, p=0.0022). The moths showed no significant preference/avoidance in treatments with other acids (Fig. 7C). The subsequent dose-response experiment showed that adults of both sexes were significantly less attracted to the 10% honey solution with octanoic acid above 1 μg. No significant difference was observed between the control and the dose of 100 ng octanoic acid in either sex (female: χ^2^=0.1538, d.f.=1, p=0.6949, male: χ^2^=0.36, d.f.=1, p=0.5485) (Fig. 7D).

## Discussion

IRs have been well studied in *Drosophila* and in some mosquito species, whereas the function of this gene family remains poorly understood in other insects including the Lepidoptera. Through phylogenetic analysis of lepidopteran IRs, several large receptor lineage expansions are observed (Liu et al., 2018), among which the IR75p and IR75q lineages are the most salient due to their close relationship with acid-sensing IRs and enriched expression in adult antennae. The tandem duplications in these expansions were considered to be associated with the discrimination of structurally similar chemicals in high-resolution (Rytz et al., 2013; van Schooten et al., 2016). Here we present the first comprehensive investigation of the ligand specificities of IRs from the IR75p/q expansions.

We identified a set of 26 AsegIRs from the antennal and ovipositor transcriptomes of *A. segetum*. Based on the number of IRs identified in the antennal transcriptomes of other moth species and *Drosophila* (e.g., 17 in *Spodoptera littoralis*, 12 in *Helicoverpa armigera* and 19 in *D. melanogaster*) (Poivet et al., 2013; Liu et al., 2012; Menuz et al., 2014), our dataset appears to include most of the IRs expected to be expressed in the antennae and ovipositor of *A. segetum*. This number is, however, lower compared to those identified from the genomes of the other species (e.g., 45 in *S. litura*, 39 in *H. armigera* and 66 in *D. melanogaster*; Croset et al., 2010; Zhu et al., 2018a; Liu et al., 2018), which include the IRs expressed in the other body parts and pseudogenes. From the phylogenetic analysis, the IR75p/q expansions together with IR1.1 and 1.2 clades were grouped into a large cluster in which some receptors including IR31a, IR64a and IR75a,b,c have been reported as acid-sensors in *D. melanogaster*; IR64a.1 and IR64a.2 in the parasitoid wasp *M. mediator*; and IR75k in the mosquito *A. gambiae* (Abuin et al., 2011; Silbering et al., 2011; Rytz et al., 2013; Ai et al., 2013; Pitts et al., 2017; Shan et al., 2019). This suggested that the receptors within the IR75p/q expansions may also be involved in acid detection.

Based on the hypothesis of an “acid-sensing” cluster of receptors, all five AsegIRs within the IR75p/q expansions were functionally investigated in *Xenopus* oocytes. We characterized two functional IRs, with AsegIR75q.1 being specific for octanoic acid, and AsegIR75p.1 responding relatively broadly to C_5_-C_7_ straight chain acids and aldehydes, as well as unsaturated C_6_ alcohols. On top of this *in vitro* test at the receptor protein level, our *in vivo* GC-EAD experiments showed that both sexes of *A. segetum* were able to detect C_6_ to C_10_ fatty acids, with no significant difference in the response magnitudes. We further explored the behavioral significance of the responses to these fatty acids and found that only octanoic acid had a significant repellent effect on female and male *A. segetum* moths, indicating that AsegIR75q.1 might mediate the behavioral repellency of octanoic acid.

Fatty acids are common in nature and are normal constituents of vertebrate skin (Mullens et al., 2009; Venter et al., 2011). It has been shown in a number of studies that the medium-chain fatty acids, in particular C_8_-C_10_, are highly repellent or toxic to many arthropods, including flies, mosquitos, ticks, ants, midges, and bees to name a few (Legal and Plawecki, 1995; Legal et al., 1999; Mullens et al., 2009; Venter et al., 2011; Zhu et al., 2018b; Ramadan et al., 2020). For instance, octanoic acid and hexanoic acid are highly abundant in the fruit of *Morinda citrifolia* and function as secondary defence compounds which are lethal or toxic to most drosophilids, with the exception of *D. sechellia* which has evolved physiological adaptations to specialize on the “toxic” Morinda fruit (Matsuo et al., 2007; Andrade et al., 2017). Also, the C_6_-C_10_ saturated fatty acids act as negative signals for ovipositional responses of female mosquitos (Skinner et al., 1970; Hwang et al., 1982). In this study, we show that octanoic acid is also a repellent for a lepidopteran species, using a ‘reverse chemical ecology’ approach. IR75q.1 is well conserved across the Lepidoptera, and further studies are needed to address the question of whether the repellency of octanoic acid is conserved in lepidopteran insects, or if it has a specific ecological importance to *A. segetum*. The biological significance of the response of AsegIR75p.1 to hexanoic acid for *Agrotis* moths is still elusive as this compound was behaviorally inactive in our assay. However, a recent study suggested that hexanoic acid deterred female hawkmoths (*M. sexta*) from ovipositing on plants (Zhang et al., 2019).

We also explored the antennal distribution patterns of AsegIR75p/q and the co-receptor AsegIR8a, which showed signals from hair-like (basiconic or trichoid) sensilla, but not from coeloconic sensilla, on female and male antennae of *A. segetum.* In *D. melanogaster*, ten antennal IRs (IR31a, IR41a, IR75a, IR75b, IR75c, IR75d, IR76a, IR76b, IR84a and IR92a) are expressed in subsets of coeloconic sensilla on the antennae. The other four antennal IRs are not expressed in coeloconic sensilla but found in neurons of the arista (IR21a) and sacculus (IR40a, IR93a and IR64a) together with their co-receptors (Benton et al., 2009; Rytz et al., 2013; Ai et al., 2013). The two co-receptors, IR8a and IR25a, are broadly expressed in coeloconic sensilla in *D. melanogaster*; IR8a is also expressed in sacculus neurons while the IR25a is additionally detected, but more weakly so, in basiconic and trichoid sensilla as well as aristal and sacculus neurons (Benton et al., 2009; Rytz et al., 2013). Electrophysiological recordings from *D. melanogaster* showed that the neurons housed in coeloconic sensilla detect acids, ammonia and humidity, which indirectly suggests that the IRs define the response profiles of the coeloconic neurons (Yao et al., 2005; Benton et al., 2009). In the desert locust *Schistocerca gregaria*, IR25a and IR8a were shown to be expressed in coeloconic sensilla (Guo et al., 2014). Beyond these two species, no additional evidence suggest that IRs are generally expressed in coeloconic sensilla. However, accumulating evidence indicates that in some insects, IRs might be expressed in non-coeloconic sensilla. In the mosquitos, trichoid sensilla and grooved-peg sensilla both responded to amines and acids (Qiu et al., 2006) and it was speculated that the responses were underlined by IRs expressed in these sensilla (Liu et al., 2010; Rytz et al., 2013). In a basal insect species, *Lepismachilis y-signata* (Archaeognatha), in which IRs are the main olfactory receptors detecting different chemical classes of odors, IRs are expressed in OSNs underneath basiconic sensilla and no coeloconic sensilla were observed on the antennae (Missbach et al., 2014). It was suggested that the evolutionary origin of IRs might have been associated with basiconic sensilla and then their expression switched to coeloconic sensilla after the neopteran insects originated (Missbach et al., 2014).

Based on the limited information regarding the sensory physiology of coeloconic sensilla in Lepidoptera, it seems that the response profiles of coeloconic, basiconic and trichoid sensilla partly overlap despite their striking morphological differences. In *B. mori*, although some coeloconic sensilla were specialized towards C_4_-C_9_ carboxylic acids, coeloconic sensilla responding to aldehydes and alcohols were also abundant; both coeloconic and basiconic sensilla responded to hexanoic acid in *B. mori* and *M. sexta* (Pophof, 1997; Zhang et al., 2019). In addition, no thermo- or hygroreceptors were found in coeloconic sensilla of *B. mori* (Pophof, 1997). It is therefore likely that the expression of IRs is not restricted to coeloconic sensilla. Our study provides the first evidence from Lepidoptera that AsegIR75p/q and the co-receptor (AsegIR8a) are expressed in OSNs located in basiconic or trichoid sensilla, suggesting that the expression of IRs in specific sensillum types might be taxon-specific. This also suggests that the coeloconic structure is not a prerequisite for IR function in neopteran insects, at least not for acid-sensing IRs. However, further studies are needed to investigate at which taxonomic level or where in the insect phylogeny this change in expression pattern has occurred.

Gene duplications are considered an important mechanism through which organisms can acquire new genetic material for specific adaption and evolution of new functions (Crow and Wagner, 2006; Moleirinho et al., 2011). After gene duplication, the duplicated genes are less constrained by purifying selection and may start to accumulate mutations resulting in novel functions, or in most cases to accumulate loss-of-function mutations generating pseudogenes (Moleirinho et al., 2011; Andersson et al., 2015; Benton, 2015). Gene subfunctionalization can also happen which means that both pre-existing and duplicated genes distinctively but complementarily retain some of their original function (Force et al., 1999; Moleirinho et al., 2011). Our phylogenetic tree shows that two mosquito (*A. gambiae*) IRs, AgamIR75k and AgamIR75l clustered within the IR75p/q expansions and are closer to the IR75q lineage. AgamIR75k is also an acid-sensing receptor, showing primary responses to octanoic and nonanoic acids, and secondary responses to hexanoic, heptanoic, and decanoic acids (Pitts et al., 2017). The response spectrum of AgamIR75k seems to be a combination of the response profiles of AsegIR75p.1 and AsegIR75q.1, which indicates functional conservation of the IR75p/q expansions across different insect orders and also suggests that subfunctionalization has occurred after gene duplication in the IR75p/q expansions in Lepidoptera. The expression level of AsegIR75q.1 was much higher than that of AsegIR75p.1 and other IRs within the same clade, implying that the key ligand of AsegIR75q.1, octanoic acid, plays a biologically important role for *A. segetum* and needs to be detected with high-resolution. The other three receptors, AsegIR75p, AsegIR75p.2 and AsegIR75q.2 showed no response to any of the 46 tested compounds although they have higher expression levels than AsegIR75p.1, probably because the ligands of these receptors were not included in the test panel. An alternative explanation could be that nonfunctionalization occurred after the gene duplication.

### Concluding remarks

We present the first comprehensive investigation of ligand specificities of IR75p and IR75q expansions and found that these IRs responded to medium-chain fatty acids. All the C_6_-C_10_ saturated fatty acids elicited antennal responses in the *A. segetum* moths, whereas only the primary ligand of AsegIR75q.1, octanoic acid, acted as a behavioral repellent. We also provide direct evidence that the acid-sensing receptor AsegIR75q.1 and its co-receptor are expressed in basiconic or trichoid sensilla, indicating that the coeloconic structure is not a prerequisite for IR function in neopteran insects. Our study expands the acid-sensing IR cluster in insects and we propose that gene subfunctionalization may have played an important role after gene duplication events during olfactory specialization in the IR75p/q expansions in Lepidoptera.

## Materials and Methods

### Chemicals

A set of 46 odorants from different chemical classes were tested in this study, including acids, aldehydes and alcohols. Most of them are common plant volatiles and can elicit responses of neurons housed in coeloconic sensilla in *Drosophila* and *B. mori* (Table S1) (Pophof, 1997; Yao et al., 2005). For oocyte recordings, stock solutions were prepared by dissolving each compound to 100 mM in dimethyl sulfoxide (DMSO), which were stored at −20 °C. Before each experiment, the stock solution was diluted to indicated concentration in Ringer’s buffer (96 mM NaCl, 2 mM KCl, 5 mM MgCl_2_, 0.8 mM CaCl_2_, 5 mM HEPES, pH 7.6) with the final stimuli containing 0.1% DMSO. Ringer’s buffer containing 0.1% DMSO was used as negative control.

### Insects

The *A. segetum* used in the experiments were from the continuous culture at the Pheromone Group, Department of Biology, Lund University. Larvae were reared on an artificial bean-based diet, under a 16L:8D photoperiod, 50% relative humidity and a temperature of 25 °C. Pupae were sexed and females and males kept separately, and emerged adults were fed with 10% honey solution. Individuals used in all experiments were unmated.

### RNA extraction, transcriptome sequencing, assembly and annotation

Male and female antennae, and ovipositors were collected from 3-day old virgin *A. segetum* under a stereomicroscope. In total, four antennal samples for each sex and two ovipositor samples were prepared. Each antennal sample contained 5 pairs of antennae and each ovipositor sample contained 40 ovipositors. The collected tissues were immediately frozen on dry ice and then stored at −80°C until use. The samples were homogenized and the total RNA was extracted and purified using RNeasy Micro Kit (Qiagen, GmbH, Hilden, Germany). The quality and quantity of the DNase treated RNA samples were verified using a BioAnalyzer (Agilent).

Library preparation using the Illumina TruSeq RNA poly-A selection kit and sequencing over two lanes on an Illumina HiSeq2500 with Rapid SBS Kit v2 chemistry was performed at SciLifeLab (Stockholm, Sweden). Quality of the raw read data was assessed using FastQC (http://www.bioinformatics.babraham.ac.uk/projects/fastqc/). Trimming of adaptors, filtering of low-quality reads and removing contaminants were performed using Trimmomatic (v0.36) (Bolger et al. 2014) with a custom screening database. Clean reads were assembled using Trinity (v2.8.2) with default settings. Contigs from the Trinity output were clustered to remove redundancy using CD-HIT-EST (v 4.6.8) (Li and Godzik, 2006) with a sequence identity threshold of 0.95. The non-redundant transcript assembly was screened for putative protein-coding regions using the program TransDecoder (release 5.0.1) (http://transdecoder.github.io/) with the option of retaining hits to the PfamA domain database (Finn et al. 2014). Completeness was assessed using the BUSCO (v3.0.2b) (Simão et al., 2015) tool searching against the insecta_odb9 database of 1658 genes. Clean read data from each pooled sample was then mapped to the non-redundant transcripts using the align_and_estimate_abundance.pl script from the Trinity v2.8.2 software package using default parameters except for --est_method RSEM --aln_method bowtie2 --trinity_mode. Reads not mapping in pairs were discarded from DE analysis. The output was analyzed using the Bioconductor (Huber et al. 2015) package DESeq2 (v1.22.2) (Love et al. 2014) in R.

Initial functional annotation of assembled sequences was performed by blasting against the pooled database of nr (NCBI non-redundant protein sequences), as well as KOG/COG (Clusters of Orthologous Groups of proteins) and Swiss-Prot with threshold E-value < 1e–5. Additional blast searches were performed using the identified *A. segetum* IR genes as queries and the transcriptome as a custom database to ensure that all IR genes were discovered. The IRs were labelled following the nomenclature convention of lepidopteran IRs (Liu et al., 2018).

### Phylogenetic analysis

A phylogenetic tree was built based on the amino acid sequences of IRs from *A. segetum*, *B. mori*, *P. xylostella*, *H. armigera, D. melanogaster* and *A. gambiae*. The sequence alignment was performed using ClustalW, a built-in plugin with default settings in Geneious R9 (Biomatters Ltd. Auckland, New Zealand). To ensure the quality of the alignment and subsequent construction of the tree, short sequences (< 300 amino acids) were excluded. A maximum-likelihood tree was constructed using MEGA 7 with the WAG+G+F substitution model that was chosen using “Find best protein model” in MEGA 7. The tree was rooted with the IR8a and IR25a receptor lineage and further viewed and edited using FigTree 1.4.4.

### Quantitative PCR (qPCR)

For analysis of tissue expression patterns of IR8a and the five AsegIRs from the IR75p and IR75q lineages, tissues were dissected from 3-5 day old virgin moths, including chemosensory organs (antennae and proboscises) of both sexes, female ovipositors, as well as male abdomens, heads+thoraxes (without antennae), legs and wings. Total RNA of these tissues was extracted using the RNeasy Mini kit or RNeasy Plus Micro kit (Qiagen, GmbH, Hilden, Germany) depending on the weight of starting material and treated with DNase. First-strand cDNAs were synthesized from 250 ng total RNA with the SuperScript™ IV First-Strand Synthesis System (Invitrogen, Carlsbad, CA, USA) following the manufacturer’s instruction. Gene expression patterns were examined by qPCR on a Stratagene Mx3005P Real-Time PCR System (Agilent technologies), with GADPH and RPS3 as reference genes. Primers used for qPCR are shown in Table S2 and the primer efficiencies were validated by standard curves with 10X serial dilutions of the templates. Three biological replicates were performed in separate plates, with three technical replicates on each plate. The reaction was carried out in a 20 μL system containing 1 μL cDNA template, 10 μL Power SYBR Green PCR 2X Master Mix (Life Technologies), and 0.3 μM of each primer. The thermal cycling parameters consisted of an initial denaturation step at 95 °C for 10 min, followed by 45 cycles of 95 °C for 15 s and 60 °C for 1 min. No-template controls were run in parallel for each primer pair. A subsequent melting curve analysis was performed to ensure the primer specificity. The relative expression level of each gene to the average of reference genes was calculated by the comparative CT method (Schmittgen and Livak, 2008).

### Scanning electron microscopy (SEM)

The antennae from 2-3 day old adults were dissected and airdried. The dried samples were carefully mounted on SEM stubs and then sputter-coated with gold (Cesington 108 auto, 45 seconds, 20 mA). The preparations were examined by a scanning electron microscope (SEM; Hitachu SU3500) at 5 kV in the Department of Biology, Lund University.

### *Whole mount fluorescence* in situ *hybridization (WM-FISH)*

Whole mount fluorescence *in situ* hybridization was performed to monitor the antennal expression pattern of the IR8a and genes in IR75p/q expansions on both female and male *A. segetum* antennae (Karner et al., 2015; Zhang et al., 2016). Digoxigenin (DIG)- and biotin-labelled antisense probes of target IRs were transcribed from linearized pCS2+ vectors with the coding region of the corresponding receptors using the T7 RNA transcription kit (Roche, Mannheim, Germany). Labelled probes were subsequently fragmented to an average length of about 800 bp by incubation in sodium carbonate buffer (80 mM NaHCO_3_, 120 mM Na_2_CO_3_, pH 10.2). DIG-labelled probes were visualized using anti-DIG AP-conjugated antibodies and the Vector Red Alkaline Phosphatase (AP) Substrate Kit (Vector Laboratories, Burlingame, CA, US); biotin-labelled probes were visualized by streptavidin-horseradish peroxidase (HRP) conjugates with Cyanine 5 Plus Amplification Reagent in a TSA kit as the substrate (PerkinElmer, Boston, MA, USA). Sense probes labelled with DIG and biotin by SP6 RNA transcription kit were used as control. Antennae were dissected from the head and cut into pieces of about 4-5 mm length and then transferred to fixation solution (4% paraformaldehyde in 0.1 M Na_2_CO_3_, pH 9.5, 0.03% Triton X-100). The antennae were gently squeezed by fine tweezers to improve tissue penetration. The following fixation, hybridization, blocking, antibody and substrate incubation steps were performed according to previously described protocols (Karner et al., 2015). Finally, the tissues were mounted in Mowiol mounting medium. A confocal microscope (Leica SP8 DLS, Germany) was used to analyze the sections (Zhang et al. 2016).

### *Functional characterization in* Xenopus *oocytes*

First strand cDNA of female antennae was synthesized from 500 ng total RNA using SuperScript™ IV First-Strand Synthesis System (Invitrogen, Carlsbad, CA, USA) following the manufacturer’s instructions. Primers containing Kozak sequence (‘GCCACC’) and restriction sites (Table S3) were designed to amplify the full-length IR genes using Platinum Pfu Polymerase (Thermo Fisher Scientific). The amplified genes were then sub-cloned into the pCS2+ expression vectors. All DNA sequences were verified by Sanger sequencing, using a capillary 3130xL Genetic Analyzer (Thermo Fisher Scientific, Waltham, MA, USA) at the sequencing facility at the Department of Biology, Lund University. The sequences of the co-receptor AsegIR8a and genes in the IR75p/q clade that were cloned and functionally tested (AsegIR75p, AsegIR75p.1, AsegIR75p.2, AsegIR75q.1 and AsegIR75q.2) have been deposited in GenBank under the accession numbers MW659848-MW659853. Large quantities of purified plasmids containing AsegIRs and AsegIR8a with verified sequences were obtained using the PureLinkTM HiPure Plasmid Filter Midiprep Kit (Thermo Fisher Scientific). cRNAs of AsegIRs and AsegIR8a were synthesized from NotI (Promega) linearized recombinant pCS2+ plasmids using the mMESSAGE mMACHINE SP6 transcription kit (Thermo Fisher Scientific).

Each of the five AsegIRs from the IR75p and IR75q lineages were co-expressed with the putative co-receptor AsegIR8a in *Xenopus* oocytes, and two-electrode voltage clamp recordings were performed following previously described protocols (Zhang et al., 2010; Zhang and Löfstedt, 2013; Hou et al., 2020). In brief, oocytes were collected from ovarian tissue of female *X. laevis* (frogs purchased from European *Xenopus* Resource Centre, University of Portsmouth, UK) and treated with 1.5 mg/mL collagenase (Sigma-Aldrich Co., St. Louis, MO, USA) in Ca^2+^ free Oocyte Ringer 2 buffer (containing 82.5 mM NaCl, 2 mM KCl, 1 mM MgCl_2_, 5 mM HEPES, pH 7.5) at room temperature for 15-18 minutes. Mature healthy oocytes (stage V-VII) were isolated and co-injected with 50 ng IR75 cRNA and 50 ng IR8a cRNA and incubated for 4-7 days at 18 °C in Ringer’s buffer containing sodium pyruvate (550 mg/L) and gentamicin (100 mg/L). Two-electrode voltage clamp recordings were performed to measure the ligand-induced whole-cell inward currents from oocytes in good condition at a holding potential of −80 mV, with a TEC-03BF signal amplifier. The test compounds and Ringer’s buffer were applied to the oocytes through a computer-controlled perfusion system with exposure to stimuli at a rate of 2 mL/min for 20 s and extensive washing in Ringer’s buffer at a rate of 4 mL/min between stimulations. Data were collected and analyzed by Cellworks software (npi electronic GmbH, Tamm, Germany).

### GC-EAD

The antennal electrophysiological responses of male and female *A. segetum* to the synthetic mixture of C_6_-C_10_ straight chain acids were recorded on an Agilent 7890 gas chromatograph equipped with a flame ionization detector (FID) (Agilent Technologies, USA) and an electroantennographic detector (EAD) (Syntech, Germany). An Agilent J&W HP-5 column (30 m × 0.25 mm i.d., 0.25 μm film thickness; Agilent Technologies, USA) was used in the GC, where the inlet temperature was set at 250 °C, the transfer line was heated at 255 °C and the detector was set at 280 °C. Hydrogen was used as the carrier gas at a constant flow of 1.8 mL/min, and a 1:1 division of the GC effluent was directed to the FID and EAD, respectively. A PRG-2 EAG (10x gain) probe (Syntech, Germany) was used in the recording. After cutting off the tips, both antennae associated with the head of a 1-2 d old adult were mounted on the probe using conductive gel (Blågel, CefarCompex, Sweden). Charcoal-filtered and humidified air passed over the antennal preparation. The GC oven was programmed from initial hold at 100 °C for 2 min, then increased to 160 °C at a rate of 10 °C/min and then held for 2 min at the final temperature. For each recording, a synthetic mixture containing 500 ng of each acid was injected in split mode (ratio of 40:1). Data were collected with the software GC-EAD Pro Version 4.1 (Syntech, Germany). Statistical analysis between stimuli within each group was performed by one-way-ANOVA followed by an LSD test at 0.05 level. Differences between males and females responding to the same stimulus was analyzed by Student’s t-Test at p < 0.05 level. N=6 for female, N=4 for male.

### Two-choice Y-tube experiments

A Y-tube olfactometer was used to investigate the behavioral responses to acids. The olfactometer consisted of a 14 cm main arm with an insect introduction chamber attached and two 14 cm long arms with an interior angle of 120° for the treatment and control odors. The inner diameter of the Y-tube was 1.6 cm. The tests were always conducted in a dark room under red light illuminating the olfactometer from above (~26 Lux). Tests were performed during the first two hours of the scotophase at the following experimental conditions: 24°C air temperature, 55% relative humidity and 0.8 m/s airflow.

To test whether the honey solution is attractive to the adult moths, two cotton balls (loaded 1mL 10% honey solution with 10 μL acetone vs 1 mL water with 10 μL acetone) was placed at inside of each Y-arm. Individual adults of *A. segetum* were released from the induce chamber at the end of main arm (29 females and 22 males). Another control experiment (cotton balls loaded 1 mL 10% honey solution with 10 μL acetone were placed at both sides of Y-tube) were performed to check whether there is a side-bias.

In this bioassay, five fatty acids were tested individually at the dose of 100 μg, including hexanoic acid, heptanoic acid, octanoic acid, nonanoic acid and decanoic acid (N ≥ 26 for each acid and each sex). Stock solutions of these fatty acids were prepared by diluting the neat compounds to 10 mg/mL in acetone. Before each experiment, 10 μL stock solution was mixed in 1 mL 10% honey solution, then the mixture was loaded on a cotton ball. 10% honey solution with 10 μL acetone was used as control. The cotton balls were left for 10 min for the evaporation of acetone before tests and placed at inside of the ends of the Y-arms. The Y-tube and cotton balls were renewed after every 10 tests. 16-20 h starved naïve individual males/females, 2-5 days old, were used in the experiment. After the screening-assay, octanoic acid was tested at a series of doses in 10-fold increments from 100 ng to 100 μg. Insects were considered to have made a choice only when they entered either of the arms and fed on the cotton within 5 min. By contrast, during the observation period (5 min), if the insect kept moving back and forth or stayed in the main arm without movement, it was considered as no choice had been made. The proportions of no choice for each treatment and each sex were lower than 5%, so we excluded the no choice data when we did the Chi2 test for statistical significance.

## Declarations

### Ethics approval and consent to participate

The care and use of *Xenopus laevis* frogs in this study were approved by the Swedish Board of Agriculture, and the methods were carried out in accordance with the approved guidelines.

### Consent for publication

Not applicable.

### Availability of data and materials

The datasets used and/or analysed during the current study are available from the corresponding author on reasonable request. The sequence reads have been deposited in the SRA database at NCBI (BioProject accession number PRJNA707654). The amino acid sequences of the IRs identified from *Agrotis segetum* antennal and ovipositor transcriptomes are available in Additional file 2; sequences of the co-receptor AsegIR8a and IRs in the 75p/q clade (AsegIR75p, AsegIR75p.1, AsegIR75p.2, AsegIR75q.1 and AsegIR75q.2) have been deposited in GenBank (accession numbers MW659848-MW659853).

### Authors’ contributions

XQH, DDZ and CL conceived the study. XQH, DDZ, MNA, DP and CL contributed to experimental design. XQH and DP performed transcriptomic data analysis. XQH performed the molecular work, oocyte recordings, scanning electron microscopy imaging, phylogenetic analysis and behavioural assay. DDZ performed qPCR experiment. DDZ and XQH did *in situ* hybridization. HLW performed the GC-EAD. XQH and DDZ analyzed the data. XQH drafted the manuscript with contributions from HLW and DP. DDZ revised the manuscript according to reviewers’ comments. All authors provided editorial and scientific input on the final version of the manuscript.

## Acknowledgements

We are grateful to Ola Gustafsson at the Microscopy Facility, Department of Biology, Lund University for access to the scanning electron microscope and confocal microscope. We thank Prof. Staffan Bensch’s lab for providing the qPCR facility and Erling V. Jirle for his help with the technical assistance and maintenance of the insect culture.

## Figure legends

**Additional file 1: Table S1.** Compounds used for characterization of *Agrotis segetum* ionotropic receptors (IRs), including their purities and source information. **Table S2.** Primers used in qPCR. **Table S3.** Primers used in gene cloning. **Fig. S1.** Additional representative FISH images.

