## Supplementary material for "Ionotropic receptors in the turnip moth *Agrotis segetum* respond to repellent medium-chain fatty acids": Table S1,Table S2,Table S3,Fig. S1

Table S1. Compounds used for characterization of *Agrotis segetum* ionotropic receptors (IRs), including their purities and source information.

| **Class** | **Compounds** | **Purity (%)*** | **Source*** |
| --- | --- | --- | --- |
| Acid | Acetic acid | 99% | Aldrich |
|  | Propanoic acid | >99% | G.B. |
|  | Butyric acid | 99% | ICN |
|  | Pentanoic acid | >99% | Aldrich |
|  | Hexanoic acid | >96% | G.B. |
|  | Heptanoic acid | >96% | ICN |
|  | Octanoic acid | 99% | ICN |
|  | Nonanoic acid | >95% | G.B. |
|  | Decanoic acid | >96% | G.B. |
|  | Undecanoic acd | >96% | G.B. |
|  | Dodecanoic acid | 95% | G.B. |
|  | Isobutyric acid | >96% | G.B. |
|  | 2-Methylbutyric acid | >96% | G.B. |
|  | 3-Methylbutyric acid | >96% | G.B. |
|  | 3-Methylpentanoic acid | >96% | G.B. |
| Aldehyde | Propanal | 97% | Aldrich |
|  | Butanal | 99% | Aldrich |
|  | Pentanal | 99% | Sigma |
|  | Hexanal | 96% | Sigma |
|  | Heptanal | >96% | G.B. |
|  | Octanal | 99% | Acros |
|  | Nonanal | 98% | Acros |
|  | Decanal | 98% | Sigma |
|  | Undecanal | 97% | Acros |
|  | Benzaldehyde | >98% | Sigma |
|  | 3-Methylbutanal | >96% | G.B. |
|  | *E*-2-Hexenal | 98% | Aldrich |
|  | Phenylacetaldehyde | 95% | ICN |
| Alcohol | Propanol | 99% | Fluka |
|  | Butanol | 99% | Aldrich |
|  | Pentanol | 99% | Sigma |
|  | Hexanol | >93% | Sigma |
|  | Heptanol | 96% | Ekokem |
|  | Octanol | >95% | G.B. |
|  | Nonanol | >90 | G.B |
|  | Decanol | 99% | ICN |
|  | Undecanol | 99% | Aldrich |
|  | Isobutanol | 99% | Lancaster |
|  | (±)-Linalool | 97% | Fluka |
|  | Benzyl alcohol | 99% | Aldrich |
|  | (*S*)-2-Heptanol | >93(96% ee) | W.F. |
|  | (*R*)*-2-Heptanol* | >92(96% ee) | W.F. |
|  | *Z-2-Hexenol* | 95% | Aldrich |
|  | *E*-2-Hexenol | 96% | Aldrich |
|  | *Z*-3-Hexenol | 98% | Aldrich |
|  | *E*-3-Hexenol | 98% | Aldrich |

G.B. = Gift from Prof. Gunnar Bergström, Chemical Ecology, Gothenburg. W.F. = Gift from Prof. Wittko Francke, University of Hamburg, Germany. ICN = ICN Biochemicals Inc.

Table S2. Primers used in qPCR.

| Genes | Primer sequence (5’-3’) |
| --- | --- |
| AsegIR8a_F | CGGAGAGAGAGGAAGTCATAGA |
| AsegIR8a_R | AGCGAGGTCTTTCTGATTGG |
| AsegIR75p_F | GAGGTTGCATCTACCAAGAGAG |
| AsegIR75p_R | TACCGACCAGTCGCATTTATC |
| AsegIR75p.1_F | GGATGTCCAGGGACTGATAGA |
| AsegIR75p.1_R | GTTTAGTAGGAGCCAGCGATATG |
| AsegIR75p.2_F | TCACCATGTCCACCGTTATTC |
| AsegIR75p.2_R | ATCTTTGCGATGGAGTCGTATT |
| AsegIR75q.1_F | ACCATCCCGACACCAATTAC |
| AsegIR75q.1_R | CTCATCCAGACACCAGTTGAA |
| AsegIR75q.2_F | GATCTGGACTGCCCTGATATTG |
| AsegIR75q.2_R | TCTGAAGAAGACCCATCGAAAC |
| AsegRPS3-F | AAGTTCGTAGACGGCCTCAT |
| AsegRPS3-R | TTCCGAGTACTCCTTGCCTTAG |
| GADPH_F | CTTAGAAGGTGGTGCCAAGAA |
| GADPH_R | GGAGATGACCTTGTAAGATGGG |

F: sense primer; R: antisense primer.

Table S3. Primers used in gene cloning.

| Genes | Primer sequence (5’-3’) |
| --- | --- |
| AsegIR8a_F | *CCG*CTCGAG**GCCACC**ATGTCATTTTATTATTTGTTTTTGCTAATTTTTCTTATC |
| AsegIR8a_R | *GC*TCTAGACTAAGGCCTATAATCTTTAGGAAAAACACTAA |
| AsegIR75p_F | *CG*GGATCC**GCCACC**ATGTTAGTCATGAATATAATTTCGTTCACATTTTTC |
| AsegIR75p_R | *GC*TCTAGATTAACGTACGGTCTTTATCATTTGCAC |
| AsegIR75p.1_F | *GA*AGGCCT**GCCACC**ATGATCGTGAAAACATTTCTTTTGATAAGTTC |
| AsegIR75p.1_R | *GC*TCTAGATCAGTTCCGGAACGGGATGT |
| AsegIR75p.2_F | *CG*GAATTC**GCCACC**ATGTTGAAGATATCTAACATGACACCACT |
| AsegIR75p.2_R | *CCG*CTCGAGTCAAGCAGAGTTTGATACATCCGAA |
| AsegIR75q.1_F | *CG*GAATTC**GCCACC**ATGAAGATTTTAACATTAATCACAAATATAATGTGTTTA |
| AsegIR75q.1_R | *CCG*CTCGAGTTAAATGTTGCTTCGACTGCTCTCT |
| AsegIR75q.2_F | *CG*GGATCC**GCCACC**ATGTCAAATAGGTACCTAAAATCGTACCG |
| AsegIR75q.2_R | *CG*GAATTCTTAATGGTTACTTTTGTGTTTCATGCAGT |

F: sense primer; R: antisense primer. The underlined indicate restriction recognition sites, the italic indicate bases flanking the recognition sequences, and the bold indicate Kozak sequence.

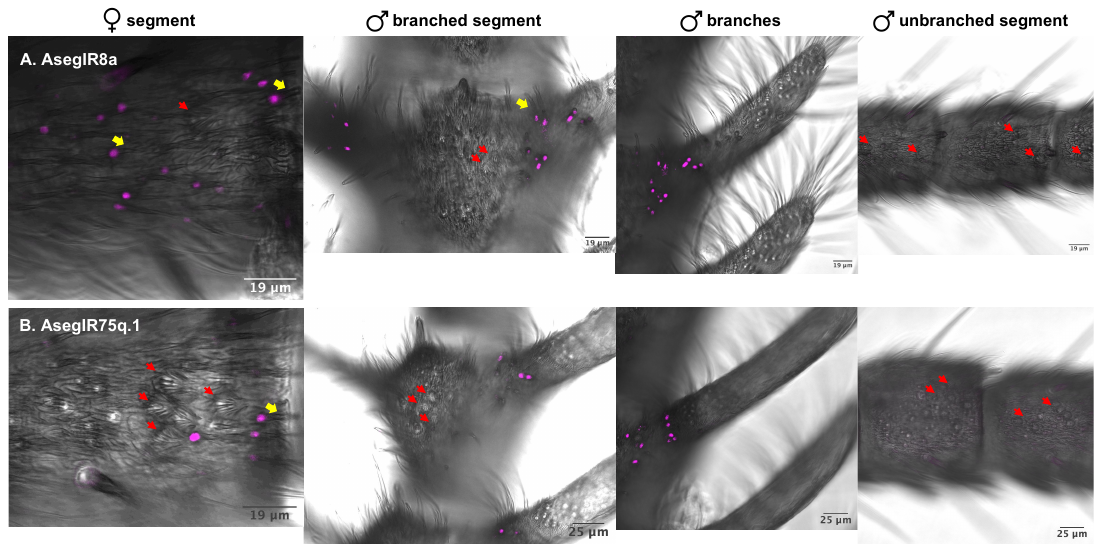

**Fig. S1.** **Additional representative FISH images.** Signals from DIG labelled AsegIR8a **(A)** or AsegIR75q.1 **(B)** probes are shown in magenta. AsegIR8a and AsegIRp/q were expressed at the branched segments of the male antennae (stem parts or bases of the branches, not on the branches), but not at the unbranched segments. Red arrows indicate the position of coeloconic sensilla and yellow arrows point to basiconic sensilla. AsegIR8a and AsegIRp/q were not expressed in coeloconic sensilla, but likely in basiconic or trichoid sensilla. The moths used in WM-FISH were 3-5 days old after eclosion and unmated.
